# Remarkable improvement in detection of readthrough downstream-of-gene transcripts by semi-extractable RNA-sequencing

**DOI:** 10.1101/2022.10.03.510745

**Authors:** Junichi Iwakiri, Kumiko Tanaka, Takeshi Chujo, Tomohiro Yamazaki, Goro Terai, Kiyoshi Asai, Tetsuro Hirose

**Affiliations:** Graduate School of Frontier Sciences, University of Tokyo, Kashiwa, 277-8562, Japan; Institute for Genetic Medicine, Hokkaido University, Sapporo 060-0815, Japan; Department of Molecular Physiology, Faculty of Life Sciences, Kumamoto University, Kumamoto 860-8556, Japan; Graduate School of Frontier Biosciences, Research Initiatives (OTRI), Osaka University, Suita 565-0871, Japan; Institute for Open and Transdisciplinary, Research Initiatives (OTRI), Osaka University, Suita 565-0871, Japan

**Keywords:** architectural RNA, DoG, hyperosmotic stress, nuclear body, readthrough transcription

## Abstract

The mammalian cell nucleus contains dozens of membrane-less nuclear bodies that play significant roles in various aspects of gene expression. Several nuclear bodies are nucleated by specific architectural noncoding RNAs (arcRNAs) acting as structural scaffolds. We have reported that a minor population of cellular RNAs exhibits an unusual semi-extractable feature upon using the conventional procedure of RNA preparation and that needle shearing or heating of cell lysates remarkably improves extraction of dozens of RNAs. Because semi-extractable RNAs, including known arcRNAs, commonly localize in nuclear bodies, this feature may be a hallmark of arcRNAs. Using the semi-extractability of RNA, we performed genome-wide screening of semi-extractable long non-coding RNAs to identify new candidate arcRNAs for arcRNA under hyperosmotic and heat stress conditions. After screening stress-inducible and semi-extractable RNAs, hundreds of readthrough downstream-of-gene (DoG) transcripts over several hundreds of kilobases, many of which were not detected among RNAs prepared by the conventional extraction procedure, were found to be stress-inducible and semi-extractable. We further characterized some of the abundant DoGs and found that stress-inducible transient extension of the 3’-UTR made DoGs semi-extractable. Furthermore, they were localized in distinct nuclear foci that were sensitive to 1,6-hexanediol. These data suggest that semi-extractable DoGs exhibit arcRNA-like features and our semi-extractable RNA-seq is a powerful tool to extensively monitor DoGs that are induced under specific physiological conditions.

## Introduction

The mammalian cell nucleus is highly organized with multiple mesoscopic membrane-less organelles, known as nuclear bodies. Nuclear bodies have been recognized as phase-separated biomolecular condensates formed through liquid–liquid phase separation and related phenomena (Mao et al. 2011; Courchaine et al. 2016; Shin and Brangwynne 2017; Alberti and Hyman 2021). Most nuclear bodies contain RNAs that associate with multiple RNA-binding proteins. Among nuclear body-localized RNAs, specific long non-coding RNAs play essential architectural roles in the construction of nuclear bodies (Chujo and Hirose 2017; Yamazaki et al. 2019). NEAT1_2 is a representative architectural RNA (arcRNA) that acts as a structural scaffold of a specific nuclear body called the paraspeckle (Chen and Carmichael 2009; Clemson et al. 2009; Sasaki et al. 2009; Sunwoo et al. 2009). NEAT1_2 sequesters more than 40 RNA-binding proteins, including FUS, which possess intrinsically disordered regions that are required to build the paraspeckle structure through phase separation (Naganuma et al. 2012; Hennig et al. 2015). Thus, arcRNAs act as the scaffold of nuclear bodies by concentrating the IDR-RNA-binding proteins to locally induce phase separation to form nuclear bodies. The expression of arcRNAs, such as NEAT1_2, HSATIII, and IGSs, is induced under various stresses, such as virus infection, thermal stress, and acidic stress, which consequently promotes the formation of cognate nuclear bodies, the paraspeckle, nuclear stress bodies, and amyloid body, respectively (Biamonti and Vourc’h 2010; Audas et al. 2016; Modic et al. 2019). arcRNAs have been identified in various eukaryotic species, suggesting that RNAs are broadly used as the scaffolds of cellular bodies in eukaryotic cells (Yamazaki et al. 2019). This raises the possibility that additional arcRNA species remain to be identified, which might be conditionally expressed under specific stresses.

We recently found that NEAT1_2 arcRNA exhibits a remarkable semi-extractable feature during the conventional RNA extraction procedure using AGPC (acid guanidinium thiocyanate-phenol-chloroform) reagent such as TRIzol or TRI reagent; syringe shearing or heating at 55°C of lysed cells in the AGPC reagent (termed as the improved method) markedly improved NEAT1_2 extraction by up to 20-fold (Chujo et al. 2017). The unique semi-extractable feature requires FUS protein which is essential for phase-separated paraspeckle assembly. This raises an intriguing possibility that the semi-extractable feature is a hallmark of arcRNAs (Chujo et al. 2017). Searching for additional RNAs exhibiting the semi-extractable feature by RNA-sequencing (semi-extractable RNA-seq), where the numbers of RNA-seq reads were compared between conventionally extracted RNA samples and those from extracted by the improved method, revealed the existence of at least 50 semi-extractable RNAs in HeLa cells under the normal condition (Chujo et al. 2017). The relatively abundant semi-extractable RNAs were commonly localized in nuclear foci distinct from the annotated nuclear bodies, which were new arcRNA candidates (Chujo et al. 2017). This finding prompted us to search for new arcRNA candidates transcribed from the human genome by performing the semi-extractable RNA-seq of cells under particular stress conditions.

Here, we searched for novel semi-extractable RNAs in HEK293 and HAP1 cells under hyperosmotic and thermal stress conditions, respectively. In the RNA-seq data analysis, we found multiple abundant long transcripts (>100 kb) that were both stress-inducible and semi-extractable, which extended from upstream protein-coding genes. These transcripts corresponded to downstream-of-gene (DoG) transcripts produced by stress-inducible transcriptional readthrough from the upstream of protein-coding genes (Vilborg et al. 2015). By definition, DoG RNAs initiate at the promoter of a protein-coding host gene and extend as long continuous transcripts for at least 5 kb beyond the 3□ terminal polyadenylation signals of their host gene (Vilborg et al. 2015). The DoG RNAs studied to date are retained in the nucleus and likely remain associated with chromatin (Vilborg et al. 2015; Rosa-Mercado and Steitz 2022). In this study, we demonstrated that the majority of abundant stress-inducible and semi-extractable RNAs were DoG transcripts, which contained many unidentified DoGs missed by RNA-seq using conventionally extracted RNAs. Therefore, semi-extractable RNA-seq using RNA samples prepared by the improved method appeared a valid method to comprehensively identify DoG transcripts. We further characterized selected DoGs and found that they formed nuclear foci possessing a feature of membrane-less nuclear bodies formed by arcRNAs.

## Results and Discussion

### Improved detection of stress-inducible DoG transcripts by semi-extractable RNA sequencing

To identify novel arcRNA candidates induced by certain stress conditions, we applied our improved RNA purification procedure to markedly facilitate the extraction of semi-extractable RNAs (Chujo et al. 2017). As stress treatments, hyperosmotic stress (OS) and heat stress (HS) were chosen. For OS treatment, HEK293 cells were cultured in medium containing 200 mM sorbitol for 6 hours, and for HS treatment, HAP1 cells were incubated at 42°C for 2 hours. RNAs were extracted by the procedures of the conventional extraction and the improved extraction method (Fig. 1A and Materials & Methods). Using these RNA samples, RNA sequencing by NGS was carried out to identify semi-extractable transcripts whose reads were significantly increased in samples subjected to the improved extraction compared with those that underwent conventional extraction. As a result, many semi-extractable transcripts were induced upon OS, and these transcripts were extremely long and extended from upstream protein-coding genes to unannotated intergenic regions (Fig. 1B). The features of the transcripts corresponded to those of DoGs that were detected under the OS condition and possessed an extremely long 3□ extension (Vilborg et al. 2015). Thus, our data suggested that at least some DoGs exhibited semi-extractable features.

**Fig. 1.**
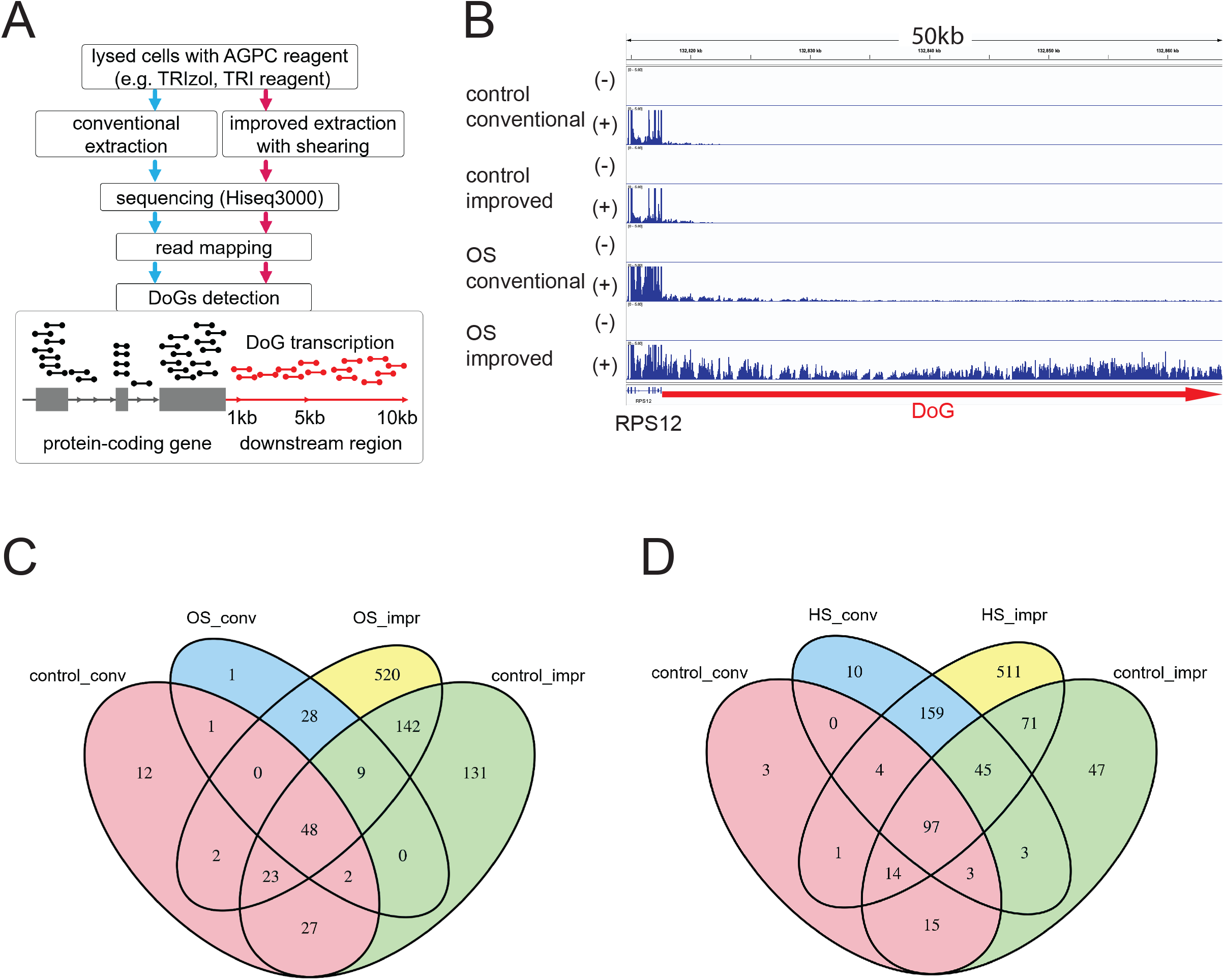
Exploratory search for stress-inducible and semi-extractable RNAs. A. Overview of semi-extractable RNA-seq and DoG detection. B. RNA-seq data of the RPS12 DoG. Trimmed mapped reads from a representative replicate of three biological replicates are shown. For each RNA preparation group, the assembled transcripts were aligned to the draft human genomic sequence (hg38). The DoG region is shown by a red arrow. The RNA samples were prepared by conventional and improved methods from unstressed control cells (control) and hyperosmotic stressed cells (OS). C, D. Venn diagrams representing four groups of DoGs based on semi-extractability and stress inducibility derived from the RNA-seq data of HEK293 cells (C) and HAP1 cells (D): (yellow) DoGs that were semi-extractable and stress-inducible, (green) DoGs that were semi-extractable, but not stress-inducible, (blue) DoGs that were stress-inducible but not semi-extractable, and (pink) DoGs that were neither stress-inducible nor semi-extractable. The numbers of DoGs in each category are also summarized in Table 1.

We focused on DoGs and identified them in our semi-extractable RNA-seq data by setting criteria applied to define the DoGs (Fig. 1A and Materials & Methods). First, semi-extractable RNA-seq data were obtained as duplicates from the RNA samples of unstressed and osmotically stressed HEK293 cells as well as from unstressed and thermal stressed HAP1 cells, which were prepared by the conventional or improved method. For HEK293 cells, 397 and 776 DoGs were identified in unstressed control cells and OS-treated cells, respectively (Supplemental Fig. 1A). For HAP1 cells, 303 and 918 DoGs were identified in unstressed control cells and HS-treated cells, respectively, most of which exhibited the semi-extractable feature (Supplemental Fig. 1B). On the basis of semi-extractability and stress inducibility, DoGs were categorized into four groups: (1) DoGs that were semi-extractable and stress-inducible, (2) DoGs that were semi-extractable, but not stress-inducible, (3) DoGs that were stress-inducible, but not semi-extractable, and (4) DoGs that were neither stress-inducible nor semi-extractable (Table 1). We identified 549 DoGs that were OS-inducible in HEK293 cells (blue and yellow parts in Fig. 1C), among which 520 (95%) were semi-extractable (yellow part in Fig. 1C). A total of 680 DoGs were detected as HS-inducible in HAP1 cells (blue and yellow parts in Fig. 1D), among which 511 (75%) DoGs were semi-extractable (yellow part in Fig. 1D). The semi-extractable features were also found in the majority of DoGs that were not stress-inducible in both HEK293 and HAP1 cells (Fig. 1C and 1D). By comparing the duplicated sequencing data, most DoGs were reproducibly found to have semi-extractable features (Supplemental Fig. 2). On the basis of these data, we propose that the majority of DoGs exhibit semi-extractable features. Therefore, semi-extractable RNA-seq using RNA samples prepared by the improved method enables extensive detection of DoGs, most of which are missed by canonical RNA-seq analyses using conventionally prepared RNA samples.

**Table 1.** Numbers of DoGs categorized into four groups based on semi-extractability and stress inducibility

| DoG group | osmotic stress | heat stress |
| --- | --- | --- |
| semi-extractable and stress-inducible | 520 | 511 |
| semi-extractable and not stress-inducible | 282 | 166 |
| stress-inducible and not semi-extractable | 29 | 169 |
| neither semi-extractable nor stress-inducible | 115 | 137 |

**Fig. 2.**
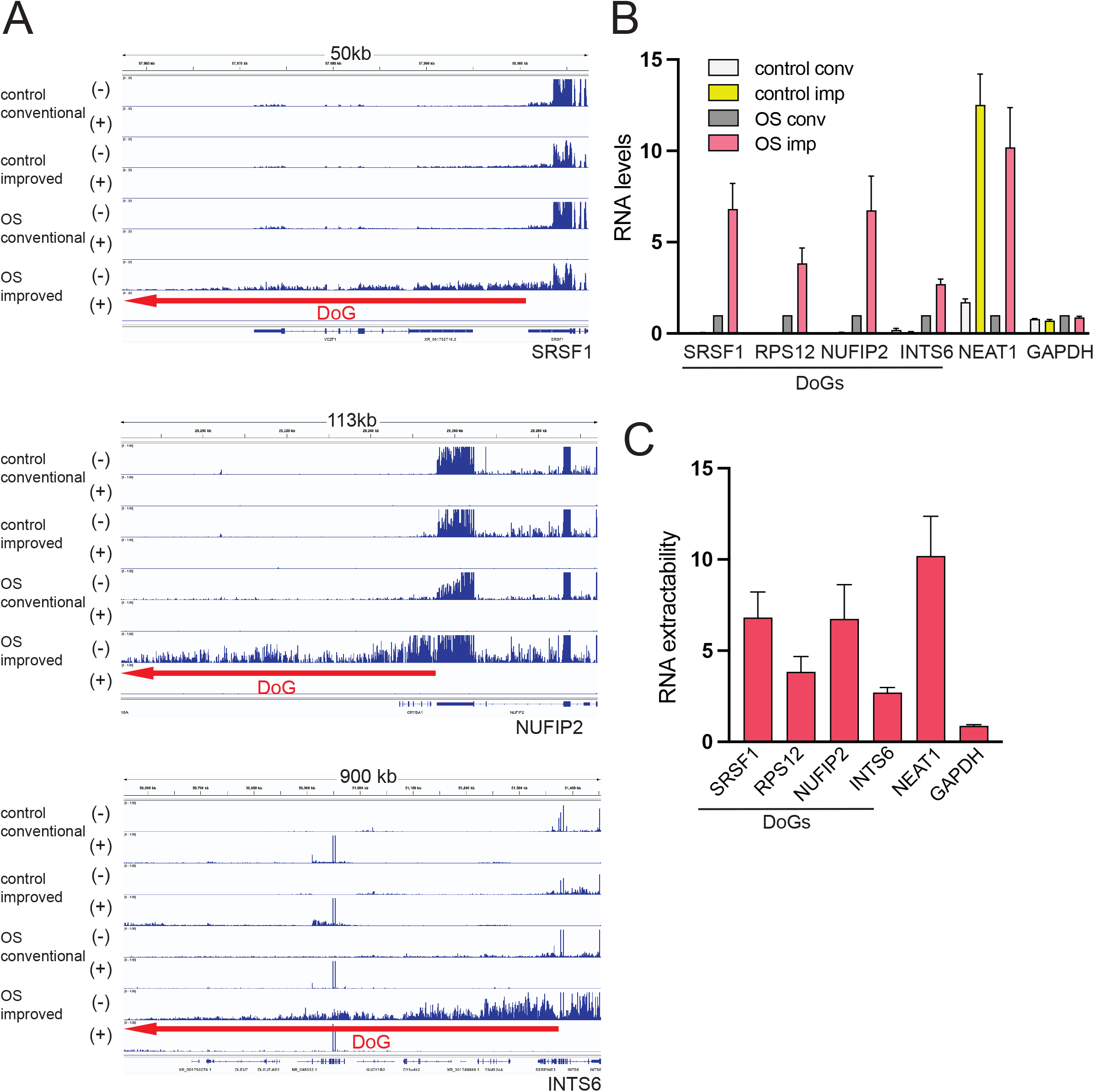
Semi-extractable feature of stress-inducible DoG transcripts. A. RNA-seq data of SRSF1, NUFIP2, and INTS6 DoGs. The DoG region is shown by a red arrow. RNA samples were prepared by conventional and improved methods from unstressed control cells (control) and hyperosmotic stressed cells (OS). B, C. RT-qPCR validation of the four DoGs in Fig. 1B and 2A (B). NEAT1 lncRNA and GAPDH mRNA were positive and negative controls of semi-extractable RNAs. RNA extractability of the transcripts in C was determined by the ratio of RNA levels extracted by improved and conventional methods. The RNA levels measured in B were normalized to the 18S rRNA level and further normalized to RNA samples extracted by the conventional method under OS. Data are shown as the mean ± SD (n = 3).

### Stress-inducible DoG transcripts exhibit a semi-extractable feature

Among the semi-extractable DoGs identified, we selected four relatively abundant DoGs for further characterization (Figs. 1A and 2). Under the OS condition, RPS12 encoding cytoplasmic ribosomal protein S12 also extended the 3’-UTR spanning >420 kb (Fig. 1B), and SRSF1 encoding Serine/arginine-rich splicing factor 1 extended the 3’-UTR spanning >150 kb downstream (Fig. 2A). Analogously, NUFIP2 and INTS6 DoGs also extended their 3’-UTRs with 400–1000 kb downstream of the original poly(A) sites (Fig. 2A). The OS-inducibility and semi-extractability of the DoGs were validated by RT-qPCR (Fig. 2B). All DoGs were detected ∼200 times and increased in the OS condition compared with those in the control condition (Fig. 2B). Simultaneously, they were 3–7-fold enriched in RNA samples prepared by the improved method compared with the conventional procedure (Fig. 2C). As a positive control of semi-extractable RNA, NEAT1 lncRNA was confirmed to exhibit a typical semi-extractable feature, but it was not OS-inducible (Fig. 2C, Supplemental Fig. 3). As a negative control, GAPDH mRNA and MALAT1 lncRNA were confirmed to be neither OS-inducible nor semi-extractable (Fig. 2C, Supplemental Fig. 3). These data confirmed that the semi-extractability was the common feature of the majority of DoGs and our semi-extractable RNA-seq was an effective method to comprehensively identify and/or monitor DoGs under specific conditions.

**Fig. 3.**
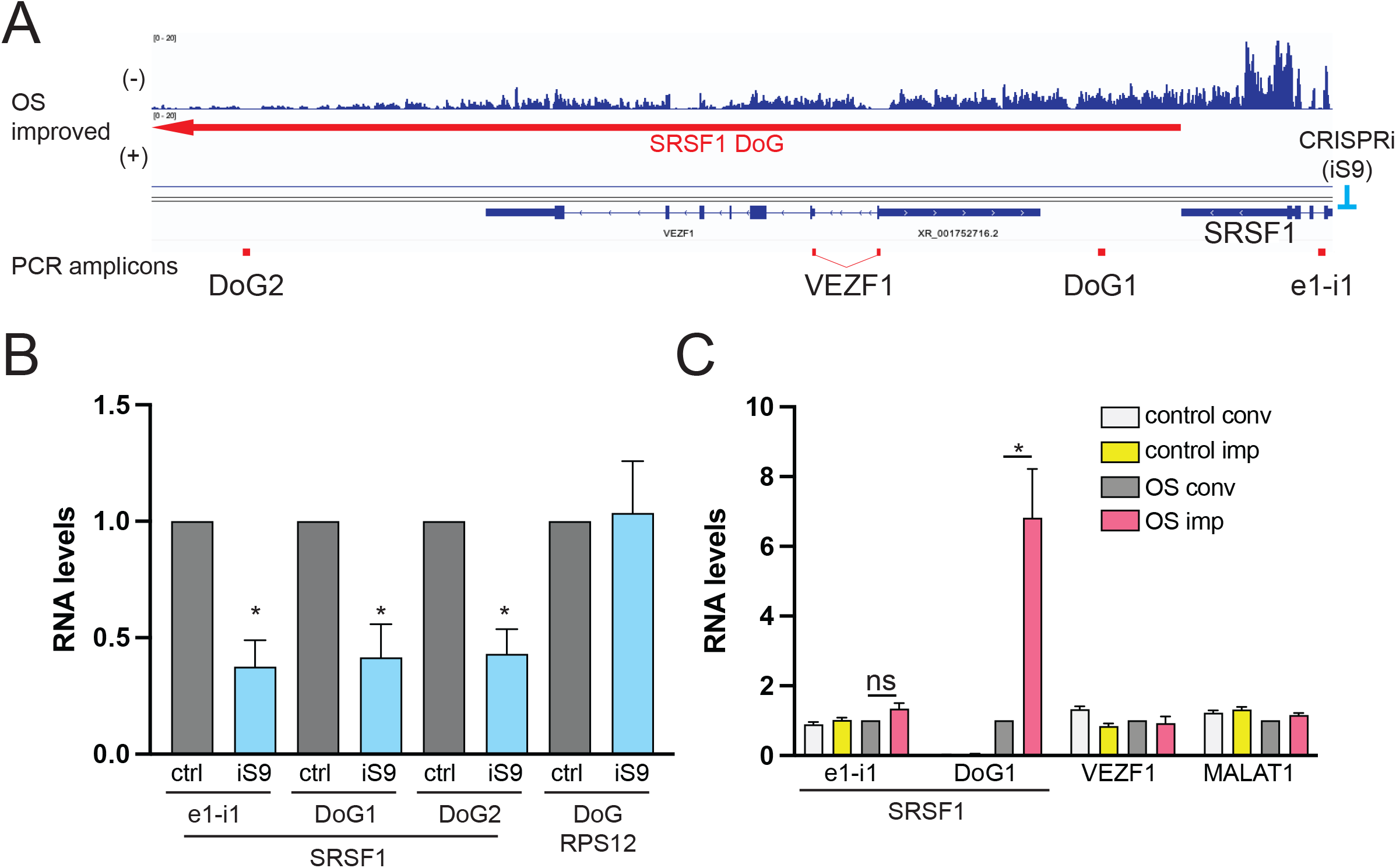
Semi-extractability is unique in 3’-UTR-extended DoG transcripts. A. Detailed map of the SRSF1 DoG region. The regions of PCR amplicons for B and C are shown below. CRISPR interference (CRISPRi) targeting the TSS of the SRSF1 gene is also shown on the right. B. CRISPRi of SRSF1 coordinately reduced the RNA levels of both the pre-mRNA and DoG of SRSF1. RNA levels in control (ctrl, gray bars) and CRISPRi cells (iS9, blue bars) quantified by RT-qPCR are plotted in the graph. PCR amplified regions in the Exon1–intron1 border (e1–i1) and two regions in DoG (DoG1 and DoG2). Control cells were transfected with pLV-GFP as the negative control. C. Semi-extractability is unique for SRSF1 DoG. Semi-extractability of SRSF1 pre-mRNA of the upstream region (e1–i1) and SRSF1 DoG (DoG1), and the spliced VEZF1 mRNA overlapped with SRSF1 DoG were quantified in four RNA samples as described in Fig. 2B. MALAT1 lncRNA was used as a negative control. The RNA levels measured in B and C were normalized to 18S rRNA levels and further normalized to control cell samples (B) and samples extracted by the conventional method under OS (C). Data are shown as the mean ± SD (n = 3). The asterisks represent p-values by Welch’s t-test (p<0.05) and ns represents not significant (p>0.05).

### A semi-extractable DoG is continuously transcribed from the upstream gene promoter

We next determined whether semi-extractable DoGs were consecutive transcripts from upstream genes and not transcripts from different conditional promoters. We employed a CRISPRi KRAB suppression system to downregulate transcription from the SRSF1 gene promoter and measured the effect on DoG expression (iS9 in Fig. 3A). CRISPRi targeting the SRSF1 gene promoter reduced the SRSF1 pre-mRNA level to 40% (e1–i1 in Fig. 3B). Concomitantly, the SRSF1 DoG level was also reduced to the almost same level detected at two positions (DoG1 and DoG2 in Fig. 3B). The CRISPRi of SRSF1 did not affect the RPS12 DoG (DoG RPS12 in Fig. 3B). This result indicated that the SRSF1 DoG was a continuous transcript from the upstream SRSF1 pre-mRNA with extended 3’-UTR. RT-PCR of unspliced SRSF1 pre-mRNA (e1–i1) did not exhibit marked semi-extractability, suggesting that only 3□ extended DoGs transcripts as a minor population of SRSF1 pre-mRNAs acquired this feature even if they were transcribed from the same TSS (e1–i1 in Fig. 3C). We also monitored the expression and the semi-extractability of VEZF1 mRNA that overlapped with SRSF1 DoGs (Fig. 3A). The spliced VEZF1 mRNA level was not affected by OS and did not exhibit the semi-extractable feature (Fig. 3C), suggesting that OS-inducible readthrough transcription poorly affected the expression of the overlapped downstream gene.

### Semi-extractable DoGs are associated with stress-inducible nuclear condensates

The semi-extractable features of RNAs are linked to their architectural roles in construction of nuclear condensates (Chujo et al. 2017). This raises the possibility that semi-extractable DoGs have analogous roles. We monitored the intracellular localization of SRSF1 and RPS12 DoGs by RNA-FISH that revealed distinct OS-dependent foci of SRSF1 and RPS12 DoGs in the nucleus, each of which never overlapped (Fig. 4A). The number of the DoG foci in a nucleus was variable between 1 and 15 for SRSF1 DoG and between 1 and 10 for RPS12 DoG. The majority of cells possessed 3 or 4 foci in a nucleus (Supplemental Fig. 4A). Recent studies have reported that DoGs are detected in close proximity to the intronic signal of neighboring genes, suggesting that DoGs are retained near their own transcription sites on a chromosome and localized in nuclear puncta (Vilborg et al. 2015; Vilborg et al. 2017). Our data supported this observation, but larger numbers of DoG foci (>10) than those of their transcription sites were present in a minor population of cells, suggesting occasional dissociation of DoG foci from their transcription sites. We also confirmed that SRSF1 and RPS12 DoG foci did not overlap with annotated nuclear bodies examined (Supplemental Fig. 4B). The DoG foci were likely analogous to the other nuclear bodies, such as the paraspeckle, which are formed around semi-extractable arcRNAs such as NEAT1_2. The 1,6-Hexanediol (1,6-HD), which is an aliphatic alcohol, interferes with multivalent hydrophobic interactions of component proteins and disintegrates phase-separated membrane-less nuclear bodies such as paraspeckles (Yamazaki et al. 2018). To further characterize the properties of DoG foci, we examined their sensitivity to 1,6-HD. After 6 hours of OS exposure, HEK293 cells were treated with 0%–8% 1,6-HD for 5 min. Treatment with >6% 1,6-HD led to the disappearance of RPS12 DoG foci, and SRSF1 DoG foci had disappeared the treatment with >8% 1,6-HD (Fig. 4B). Both DoG foci were resistant to 8% 2,5-HD as an ineffective analog (Fig. 4B). The 1,6-HD sensitivity of DoG foci was similar to that of other phase-separated nuclear bodies such as paraspeckles and Cajal bodies (Yamazaki et al. 2018), suggesting that the molecular interactions constituting DoG foci were at least in part similar to known phase-separated nuclear bodies such as paraspeckles.

**Fig. 4.**
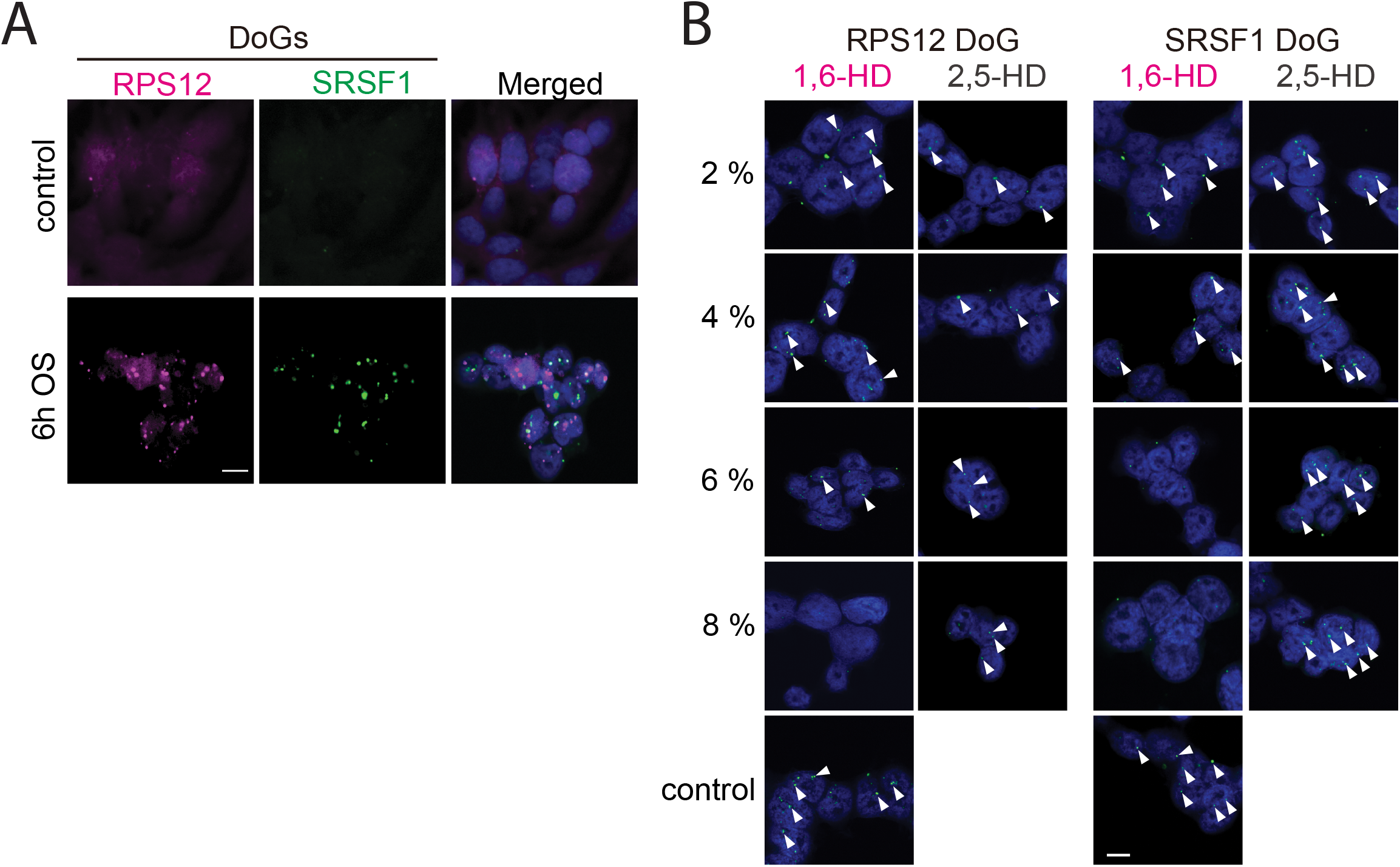
DoG transcripts are associated with OS-inducible nuclear condensates. A. Representative images of double fluorescence in situ hybridization (FISH) using RNA probes for RPS12 DoG (magenta; left) and SRSF1 DoG (green; middle). Nuclei were counterstained with DAPI (blue). Scale bars correspond to 10 μm. B. The biochemical property of DoG foci was monitored by the sensitivity of 1,6-hexanediol (1,6-HD). OS-induced DoG foci in HEK293 cells were observed upon treatment with 0%–8% 1,6-HD and 2,5-HD for 5 min. DoG foci were detected by RNA-FISH as described in A. Scale bars correspond to 10 μm.

Overall, both SRSF1 and RPS12 DoGs were relatively abundant, semi-extractable, and localized in 1,6-HD-sensitive nuclear bodies, all of which are characteristics of other arcRNAs (Chujo et al. 2017). The appearance of DoG foci correlated to DoG synthesis upon OS, suggesting that DoGs contributed to the conditional formation of nuclear bodies near their own transcription sites. The biological significance of the formation of DoG foci during OS remains to be understood. Some nuclear bodies act to sequester specific factors to regulate gene expression, raising the intriguing possibility that DoG foci also sequester specific factors for RNA processing and/or export to regulate transcriptional and post-transcriptional events during stress conditions. Finally, our semi-extractability RNA-seq was potent to extensively identify DoGs under various conditions. Further analyses of the newly identified DoGs may provide new insights into the roles of global transcriptional readthrough during specific physiological conditions and diseases.

## Materials and Methods

### Cell culture and induction of osmotic stress

HEK293 cells were obtained from the American Tissue Collection Center. The cells were cultured in Dulbecco’s modified Eagle’s medium (Nacalai Tesque) supplemented with 10% (v/v) fetal calf serum (Sigma-Aldrich) and maintained at 37°C in a humidified atmosphere with 5% CO_2_. To induce osmotic stress (OS), HEK293 cells were exposed for 6 hours to 200 mM sorbitol dissolved in culture medium. To induce heat stress (HS), HAP1 cells were exposed at 42°C for 2 hours. For aliphatic alcohol treatment, the indicated concentrations of 1,6-hexanediol or 2,5-hexanediol dissolved in culture medium were added to cells post-OS treatment, followed by incubation for 5 minutes at room temperature.

### RNA extraction

Total RNA was extracted using 1 mL TRI Reagent (MRC) per 1 × 10^6^ cells. The improved method was applied to purify semi-extractable RNA. In detail, the resulting cell lysates were passed through a 20 G needle 100 times or heated using a Thermomixer (Eppendorf) at 55°C for 10 minutes with 1000 rpm agitation prior to the conventional RNA purification procedure in accordance with the manufacturer’s protocol.

### RNA-sequencing

Global RNA expression profiles were compared between total RNA prepared by conventional or improved methods from normal HEK293 cells and cells exposed to osmotic stress for 6 h. An aliquot of 1 μg purified RNA was applied to RT-qPCR and the remaining was subjected to ribosomal RNA depletion using a Ribo-Zero Gold Kit (Epicentre). Sequencing libraries were constructed from 100 ng total RNA using a Truseq stranded mRNA Library Prep Kit (Illumina) without the poly-A selection step. Subsequently, sequencing was performed using a Hiseq3000 (Illumina) with the 36 or 50 bp single-end method.

### RNA-seq data processing and DoG detection

Low-quality bases in raw RNA-seq reads were trimmed and reads containing ambiguous bases were filtered out using cutadapt (version 2.0) (Martin 2011) with the parameters -q 30 --max-n 0 -m 30. The remaining reads were aligned to human rRNA sequences (accession ID: U13369.1 and V00589.1) to remove reads derived from rRNAs using STAR aligner (version 2.5.3a) (Dobin et al. 2013) with the parameters --outFilterMultimapNmax 100 --outReadsUnmapped Fastx. Unmapped reads were aligned to the human reference genome (hg38) with GENCODE annotation (release 27) (Frankish et al. 2019) using STAR aligner with the parameter --outFilterMultimapNmax 1. Reads mapped to regions annotated in GENCODE were counted using featureCounts (version 1.4.6) (Liao et al. 2014) with the parameters: -s 2 -t exon -g gene_id. To detect DoG transcripts of a protein-coding gene, reads mapped to the downstream region of a gene needed to be counted without miscounting reads derived from other downstream protein-coding genes. Among the protein-coding genes in the GENCODE annotation, 14,930 genes whose 10-kb downstream regions were not overlapped with any other annotated genes were selected for the subsequent DoG candidate detection. To detect DoG candidates derived from protein-coding genes, expression levels (FPKM) were calculated for their exonic region (*FPKM*_*exon*_), and 1-kb (*FPKM*_*down1kb*_), 5-kb (*FPKM*_*down5kb*_), and 10-kb (*FPKM*_*down10kb*_) downstream regions. The following criteria were used to detect DoG candidates: (a) *FPKM*_*exon*_>10, (b) *FPKM*_*down1kb*_>0.05×*FPKM*_*exon*_, (c) *FPKM*_*down5kb*_>0.05×*FPKM*_*exon*_, (d) *FPKM*_*down10kb*_>0.05×*FPKM*_*exon*_. Genes reproducibly detected as candidates in both biological replicates were nominated as DoGs. The overall data of the semi-extractable RNA-seq are associated with those under other conditions and separately reported in the other paper (Zeng et al., 2022).

### Classification of DoGs

DoGs detected in the RNA-seq data were further characterized by RNA extractability and stress inducibility. DoGs detected in samples prepared by improved RNA extraction, but not detected in samples prepared by conventional extraction, were classified as semi-extractable DoGs. The DoGs detected in samples exposed to hyperosmotic stress or heat stress, but not in the normal condition samples, were classified as stress-inducible DoGs. Finally, DoGs are categorized into following four groups: (a) semi-extractable and stress-inducible, (b) semi-extractable and not stress-inducible, (c) stress-inducible and not semi-extractable, and (d) neither semi-extractable nor stress-inducible.

### Reverse transcription-quantitative real-time PCR (RT-qPCR)

Isolated total RNA (1 μg) was treated with RQ1 RNase-Free DNase (Promega) to remove genomic DNA and then reverse transcribed using a High-Capacity cDNA reverse-transcription kit with random hexamer primers (Applied Biosystems) in accordance with the manufacturers’ instructions. For quantitative determination of RNA levels, qPCR was performed on a LightCycler® 480 instrument (Roche Diagnostics) with a reaction mixture of KAPA SYBR FAST qPCR Master Mix (Kapa Biosystems) and sequence-specific primers listed in Supplemental Table 2.

### RNA fluorescence in situ hybridization (FISH) and immunofluorescence

Cell samples plated on coverslips precoated with poly-L-lysine (SIGMA) were fixed in 4 % PFA and then processed for RNA FISH and immunofluorescence as described previously (Naganuma et al, 2012). Double fluorescence TSA was carried out using FITC-labeled doRPS12 and DIG-labeled doSRSF1 RNA probes detected with HRP-labeled anti-fluorescein and Cy3-conjugated Tyramide or HRP-labeled anti-digoxigenin antibodies with Cy5-conjugated Tyramide (PerkinElmer, Waltham, MA, USA). After probe hybridization and washing, the TSA procedure was performed in accordance with the manufacturer’s protocol, adding a 30-min treatment with 1 % of H_2_O_2_ between two TSAs to quench residual peroxidase activity from the first TSA reaction. Confocal images were acquired under a FLUOVIEW FV1000 confocal laser scanning microscope (Olympus) and a Nikon Confocal system A1Rsi (Nikon) equipped with a Plan Apo VC x60 objective lens (NA 1.40, Nikon).

### Quantification of DoG foci per nucleus

The distribution of the focus number in the nucleus was analyzed in z-stacked images obtained by cross-sections in confocal laser scanning microscopy. The captured images were analyzed to count foci using Volocity software. In both RPS12 and SRSF1 DoGs, approximately 100 nuclei with at least one focus were counted and the number of DoG foci per nucleus was displayed as a scatter dot plot. The horizontal line represents the average value with the standard deviation (SD).

### Construction of CRISPR interference (CRISPRi)-based transcription repression plasmids and transfection

pLVhU6-sgRNA hUbC-dCas9-KRAB-T2a-GFP was purchased from Addgene (Cambridge). Oligos encoding sgRNAs designed using GPP sgRNA Designer (https://portals.broadinstitute.org/gpp/public/analysis-tools/sgrna-design-crisprai) were annealed and cloned into the plasmid. HEK293 cells were washed twice with PBS and 2 × 10^6^ cells were resuspended in 100 μl of Solution V (Lonza) and then mixed with 5 μg purified sgRNA expression vector or the empty vector as a negative control before nucleofection using program Q-001 of a Nucleofector™ 2b (Lonza). To enrich transfectants, single cells were sorted 16 hours post-nucleofection using an SH800 cell sorter (SONY), and GFP-positive cells were enriched. Cells were then cultured in complete Dulbecco’s modified Eagle’s medium in two poly-L-lysine-coated wells of a 24-well plate at 37°C in a humidified atmosphere with 5% CO_2_ for 16 hours before treatment with 200 mM sorbitol for 3 hours.

## Data availability

The conventional and semi-extractable RNA-seq data have been deposited in the DDBJ Sequence Read Archive (DRA) under accession numbers DRA012807 and DRA009793.

## Acknowledgments

The authors thank the members of the Hirose and Asai laboratories for their valuable discussions. The computational analysis was partially performed on the NIG supercomputer at the ROIS National Institute of Genetics. This work was supported by JST CREST grant no. JPMJCR20E6 (to T.H.), AMED grant no. 21479280 (to T.H.), and JSPS KAKENHI grants nos. 26113002, 16H06279, 20H00448, 21H05276, and 22K19293 (to T.H.).

## Supplemental material

**Supplemental Fig. 1**. Venn diagrams of the semi-extractability of DoGs in control and OS-treated HEK293 cells (A) and those in control and HS-treated HAP1 cells (B). The numbers of DoGs in each category are shown.

**Supplemental Fig. 2**. Venn diagrams of data reproducibility in Supplemental Fig. 1.

**Supplemental Fig. 3**. RNA-seq data of NEAT1 and MALAT1 using RNA samples prepared by conventional and improved methods from control and OS-treated HEK293 cells. NEAT1 and MALAT1 lncRNAs were used as positive and negative controls of semi-extractable RNA, respectively.

**Supplemental Fig. 4**. A. Focal numbers of SRSF1 and RPS12 DoGs in a nucleus and each cell number shown by a violin plot. B. RNA-FISH of DoG RNAs and immunofluorescence staining of known markers for various nuclear bodies. The names of nuclear bodies and markers are indicated on the left. Nuclei were counterstained with DAPI (blue). Scale bars correspond to 10 μm.

**Supplemental Table 1**.

Detected DoGs with semi-extractable and stress-inducible features. Averaged expression levels (FPKM) were calculated from semi-extractable RNA-seq data of HEK293 cells under the osmotic stress condition and HAP1 cells under the thermal stress condition.

**Supplemental Table 2**. Primers used in this study

**Supplemental Table 3**. FISH probes used in this study

